# Invariant chain regulates endosomal fusion and maturation through the SNARE Vti1b

**DOI:** 10.1101/771857

**Authors:** Dominik Frei, Azzurra Margiotta, Marita Borg Distefano, Mohamed Moulefera, Lennert Janssen, Jacques Thibodeau, Jacques Neefjes, Oddmund Bakke

## Abstract

Invariant chain (Ii) is an important multifunctional player in the regulation of adaptive immune responses and is responsible for several cellular functions related to MHCI and MHCII antigen loading and antigen presentation. While regulating endosomal trafficking of MHCII and other proteins that bind to Ii, this molecule is able to influence the endosomal pathway delaying the maturation of endosomes to the late endosomal loading compartments. When expressed in cells Ii is found to increase endosomal size, but the mechanisms for this is not known. We used RNAi silencing to identify SNARE proteins controlling Ii induced increase of endosomal size and delay of the endosomal pathway. Ii was found to interact with the SNARE protein Vti1b. Vti1b localized at the contact sites of fusing Ii positive endosomes and a tailless Ii was able to relocate Vti1b to the plasma membrane. Furthermore, silencing Vti1b, abrogated the delay in endosomal maturation induced by Ii expression. In conclusion, Ii interacts with Vti1b and this interaction is fundamental for Ii-mediated alteration of the endosomal pathway. We propose that Ii, by interacting with SNAREs, in particular Vti1B in the biosynthetic pathway of antigen presenting cells, is able to assemble SNARE directed fusion partners in the early part of the endosomal pathway that lead to a slower endosomal maturation for efficient antigen processing and antigen loading.

## Introduction

Professional antigen presenting cells like dendritic cells and B cells express major histocompatibility complex class II (MHCII) and associated molecules in the MHC region and invariant chain (Ii, or CD74) located on a different chromosome (Genuardi and Saunders 1988). While MHCII is a polymorphic protein, Ii is a non-polymorphic type II transmembrane protein that forms complexes by self-trimerization and binding to MHC class II (reviewed in (Landsverk, Bakke, and Gregers 2009). Ii is mainly expressed in antigen presenting cells (APCs) and has a prominent role sorting associated molecules to the endosomal pathway (reviewed in (Schroder 2016)). Importantly, Ii, which contains two efficient leucine based endosomal sorting signals, facilitates the exit of MHC class II from the endoplasmic reticulum (ER) in antigen-presenting cells and mediates rapid transport of the MHCII/Ii complex to the endosomal pathway where the antigen is efficiently loaded (Bremnes et al. 1994; Bakke and Dobberstein 1990; Elliott et al. 1994; Bikoff et al. 1993; Neefjes and Ploegh 1992). In addition, Ii delayed transport from Rab5 positive early endosomes (EE) to late Rab7a positive endosomes (LE) (Gorvel et al. 1995). Ii also induced fusion of early endosomes creating enlarged endosomes (Pieters, Bakke, and Dobberstein 1993; Romagnoli et al. 1993). These enlarged endosomes were able to mature normally, however, with a prolonged residence time in EE (presence of EEA1 and Rab5) before the endosomes switched to late, acidic Rab7a and Lamp1 positive multivesicular endosomes and lysosomes (Stang and Bakke 1997; Landsverk et al. 2011).

Most membrane fusion events in the cell require SNARE (soluble N-ethylmaleimide-sensitive-factor (NSF) attachment protein (SNAP) receptor) proteins (reviewed in (Jahn and Scheller 2006)). All SNAREs share the conserved SNARE motif of 60-70 amino acids (Weimbs et al. 1997) and generally contain a transmembrane domains; however, some are anchored to membranes by lipid modifications. There are several types of SNARE domains: R-SNAREs contribute an arginine residue to the binding layer of the assembled complex, while Q-SNAREs have conserved glutamine residues (Fasshauer et al. 1998). Q-SNAREs are further subdivided into Qa-, Qb- and Qc-SNAREs depending on their position in the complex. One SNARE domain of each class is required to form the complex. The formation of this complex provides the driving power necessary for membrane fusion.

Interestingly, although PI3K inhibitors block homotypic fusion of early endosomes (Jones and Clague 1995; Powis et al. 1994), this is not the case for Ii-positive endosomes (Nordeng et al. 2002). This indicates that the Ii mediated fusion mechanism differ from the standard early endosomal fusion machinery. The Ii induced fusion of endosomes was, however, inhibited by NEM suggesting that SNARE molecules were involved in this process. After failed rounds of searching for Ii associated proteins explaining the trafficking effects by two-hybrid screens and coimmunoprecipitations experiments, we set out to test the involvement of every individual SNARE protein in the Ii specific endosomal fusion events.

Our siRNA screen and subsequent verification identified the Q-SNARE Vti1b as a plausible candidate. Vti1b depletion reduced Ii-induced endosomal size, indicating a functional role in this process. This was further substantiated by a co-isolation of Vti1b with Ii. Vti1b was found enriched at contact sites between Ii-positive endosomes and present at the fusion pore. As a further proof, depletion of Vti1b abrogated the effect of Ii-induced delay in endosomal maturation. These data suggest that Vti1b is an essential element in the regulation of the endosomal pathway by Ii.

## Results

### Ii regulates endosomal maturation in antigen presenting cells

In our earlier studies the Ii induced endosomal maturation delay was measured as a delayed entry from Rab5 positive to Rab7 positive endosomes (Gorvel et al. 1995) or a prolonged EE phase (Landsverk et al. 2011) in non-antigen presenting cells without the MHC domain molecules. If Ii has a similar regulatory role on endosomal maturation in cells containing the full MHC II antigen presenting machinery removal of Ii should speed up endosomal maturation. To test for this, we generated, using CRISPR-Cas9, human antigen-presenting cell lines lacking Ii (Ii KO) (see M&M). We first followed progression of the fluorescent fluid phase marker 8-hydroxypyrene-1,3,6-trisulphonate (HPTS) in human lymphoblast Raji cells either expressing the Cas9 enzyme (control) or Ii KO by measuring the time for a HPTS to colocalize with acidic lysotracker red (LTR) late endosomes. In Ii KO cells, fluid phase marker HPTS reached lysotracker red (LTR) late endosomes faster than in the control cells (Figure 1A). 1 hour after addition of HPTS the colocalization with LTR was 45% in the Ii KO cells while in the corresponding control cells, this was 30% indicating that the KO of Ii in the Raji cells causes a significant faster endosomal maturation. Meljuso cells naturally express Ii and MHCII and are a popular model system for studying antigen presentation. HPTS colocalization with LTR was followed over time in control and Ii KO MelJuSo cells for up to 200 min (Fig 1B). At this time point, in the Ii KO cells about 50% of LTR colocalized with HPTS, compared to only 30% in control cells. The entry of HPTS into late endosomal compartments was again significantly faster in Ii KO cells. This suggests that Ii regulates endosomal maturation not only in model fibroblast-like cell lines transfected with Ii, but also in antigen presenting cells that contain the full MHCII-related antigen processing and presentation machinery.

**Figure 1.**
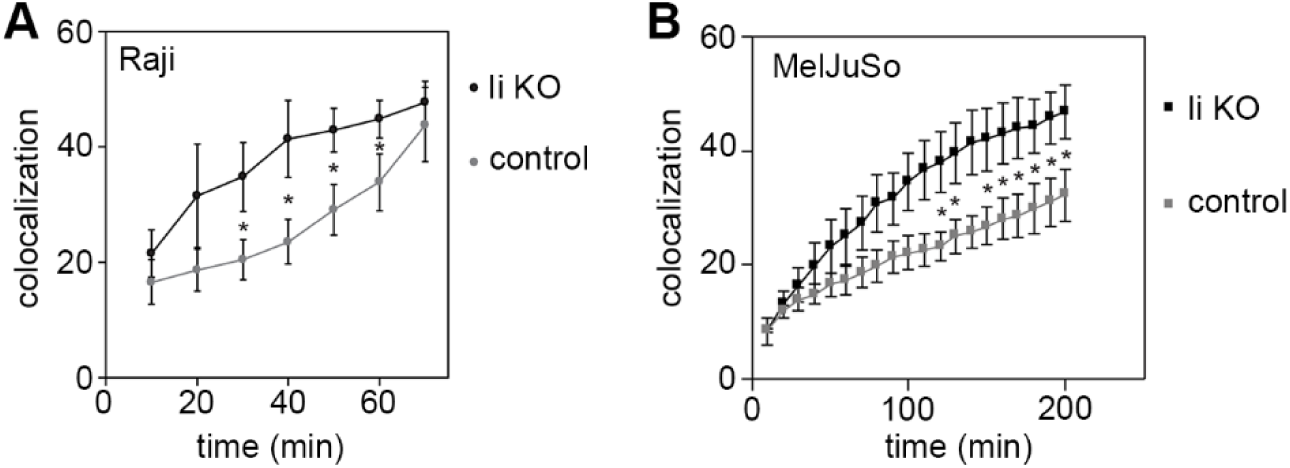
Ii KO speeds up trafficking of HPTS. A) The colocalization coefficient (% pixel overlap of LysoTracker Red with HPTS) in wt Raji cells expressing Cas9 (control) or Ii knockout (Ii KO) was analysed and is represented by line graphs for control (●) and Ii KO (●) indicating mean ± SEM of 6 independent experiments. B) Same as for A) in wt Meljuso cells (control, ●) or Ii KO (●) in showing mean ± SEM for 3 independent experiments.

### Vti1b is involved in Ii-mediated endosome enlargement

When Ii was expressed in non-antigen presenting cells, the size of the endosomal compartments strongly increased (Pieters, Bakke, and Dobberstein 1993; Romagnoli et al. 1993). In fact, there was a direct correlation between the size increase and Ii-expression levels (Nordeng et al. 2002). Both the delayed maturation and the endosomal expansion was dependent on Ii trimerization and suggested that these effects were caused by the same mechanism (Nordeng et al. 2002; Gregers et al. 2003). NEM is a chemical compound that block amongst others SNARE proteins. Since NEM inhibited Ii driven effects on the endosomal pathway, we proposed a contribution of SNAREs (Nordeng et al. 2002).

To test for SNAREs involved in Ii-mediated endosome enlargement, we silenced all individual SNAREs in the M1 fibroblast cell line, stably transfected with Ii under the control of a heavy-metal inducible promoter (M1 pMep4-Ii, (Nordeng et al. 2002)). The M1 wild type cell line is negative for Ii and the MHCII proteins but the cell line has been employed extensively to study endosomal dynamics and antigen presentation by transfecting in immune genes (Roche et al. 1993; Stang and Bakke 1997; Skjeldal et al. 2012; Sand et al. 2014; Haugen et al. 2017). Ii transport to Ii positive endosomes was visualized by adding fluorescently tagged antibody recognizing the extracellular/luminal domain of Ii, and the size of Ii-positive endosomes was analysed by confocal microscopy (Figure 2A).

**Figure 2.**
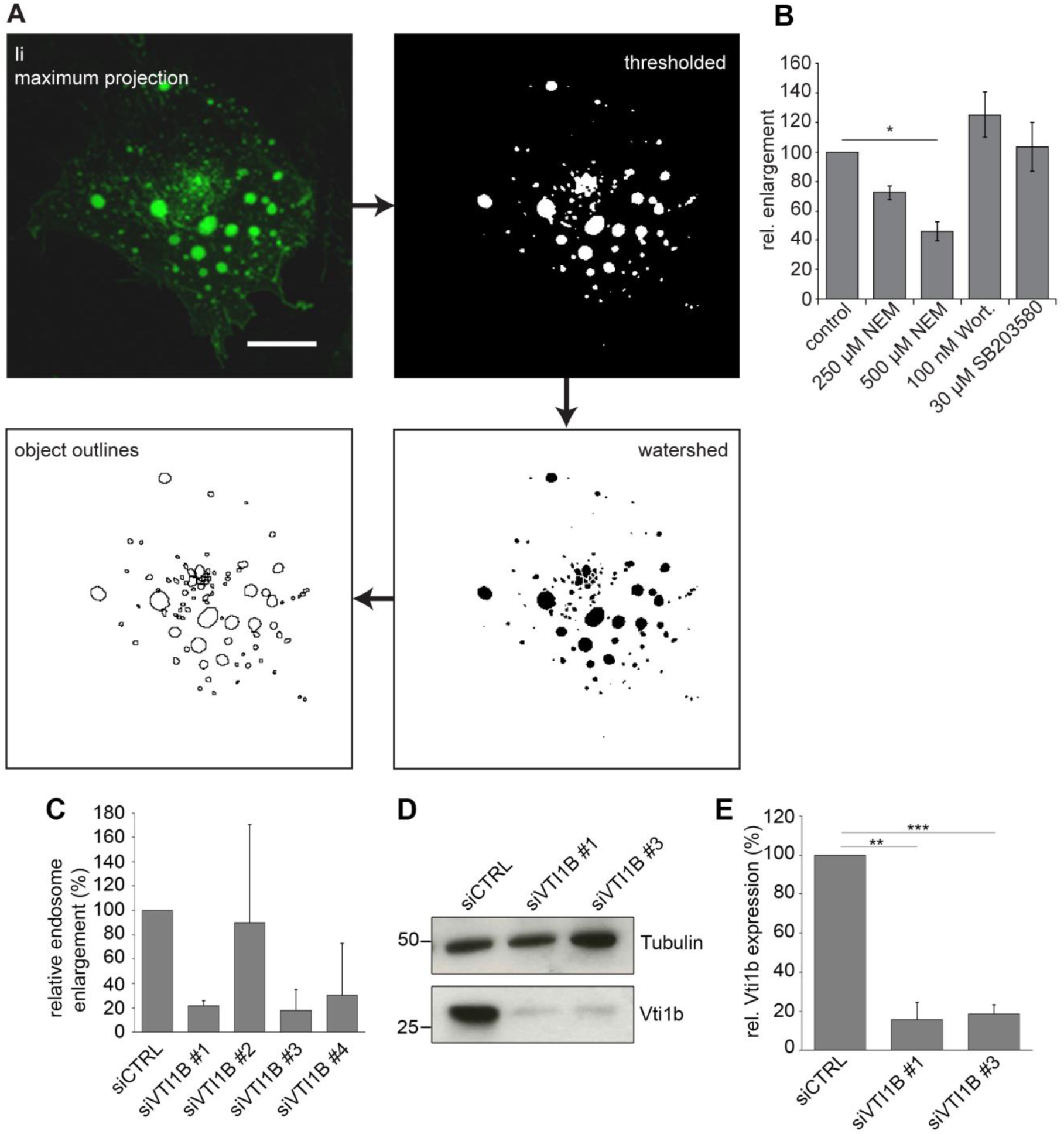
Ii-induced enlargement of endosomes is dependent of Vti1b. A) Workflow diagram of the image processing for the quantification of endosome size. Scale bar: 10 µm. C) Results of endosome quantification after treatment with different drugs affecting early endosome fusion, mean ± s.e.m. of three independent experiments. D) Quantifications results of enlarged endosomes (bigger than 3 µm in diameter) for four individual oligonucleotides targeting Vti1b compared to control siRNA (non-targeting, siCTRL) reported as relative endosome enlargement. E) M1-pMep4-Ii cells transfected with control siRNA or siRNA against Vti1b were lysed and subjected to western blot analysis using anti-Vti1b and anti-tubulin antibodies. Detection of tubulin has been used as loading control. F) Quantification of Vti1b abundance is shown. Data represent the mean ± s.e.m. of three independent experiments.

As a test for our screening system we employed some of the known inhibitors of early endosome fusion. The MAP kinase inhibitor SB203580 (Cuenda et al. 1995) was used to block Rab5 activation and Wortmannin to inhibit PI3K activity (Wymann et al. 1996). As shown earlier (Nordeng et al. 2002), wortmannin did not inhibit formation of enlarged endosomes (Fig 2B). MAP kinase activity was also not essential since SB203580 could not stop the formation of the enlarged endosomes (Fig 2B). However, N-ethylmaleimide (NEM), a general inhibitor of SNARE function, significantly reduced Ii-mediated endosome enlargement by about 50% (Fig 2B), in line with earlier results from cell free studies (Nordeng et al. 2002).

To screen for specific SNARE proteins involved in Ii-mediated endosome enlargement, inducible M1 pMep4-Ii cells were transfected with pools of siRNAs (four oligos per target gene) targeting 40 individual human SNAREs. Three days post-transfection, cells were analysed by 3D confocal microscopy to determine the degree of endosome enlargement upon Ii expression. This screen showed that silencing Vti1b counteracted the Ii induced endosome enlargement by 50%. Validation with individual siRNAs for Vti1b showed an even stronger inhibitory effect for two out of the four siRNAs (Fig 2C). Western blot analyses of cells transfected with these siRNAs confirmed a significantly reduced level of the Vti1b target protein (Fig 2D-E).

These experiments suggest that depletion of Vti1b inhibits the Ii induced formation of enlarged endosomes.

### Silencing of Vti1b specifically reduces Ii-mediated endosome enlargement

Given the effect of Vti1b in the regulation of the size of Ii-positive endosomes, we evaluated whether the depletion of this SNARE also interfered with endosomes enlarged by other means than expression of Ii. A well-known method for expanded early endosomes is to express the active mutant of Rab5 (Rab5Q79L) (Stenmark et al. 1994; Wegner et al. 2010). M1 pMep4-Ii cells were transfected with control siRNA or siRNA targeting Vti1b and subsequently either treated with CdCl_2_ to induce Ii expression, (inducible metallothionine promotor) (Fig 3A, top row) or transfected with EGFP-Rab5Q79L (Fig 3A, bottom row). Images were acquired and the size of Ii- or Rab5Q79L-positive endosomes was determined (Fig 3B). Silencing of Vti1b with either of the two siRNAs reduced Ii-mediated endosome expansion by approximately 70% (Fig 3B). On the other hand, Vti1b depletion did not affect the size of Rab5Q79L-positive endosomes, showing that Vti1b is not essential for endosome enlargement of early endosomes induced by active Rab5. To study this process further we quantified the number of endosomes per cell in the above experiment and observed that the silencing of Vti1b significantly increased the number of endosomes after Ii expression (Fig 3C). On the other hand, silencing of Vti1b had no effect on the number of endosomes after overexpression of Rab5Q79L (Fig 3C) The above measurements suggest that Vti1b effect is not a general requirement for enlarging early endosomes, but more specific for the Ii induced endosomal expansion.

**Figure 3.**
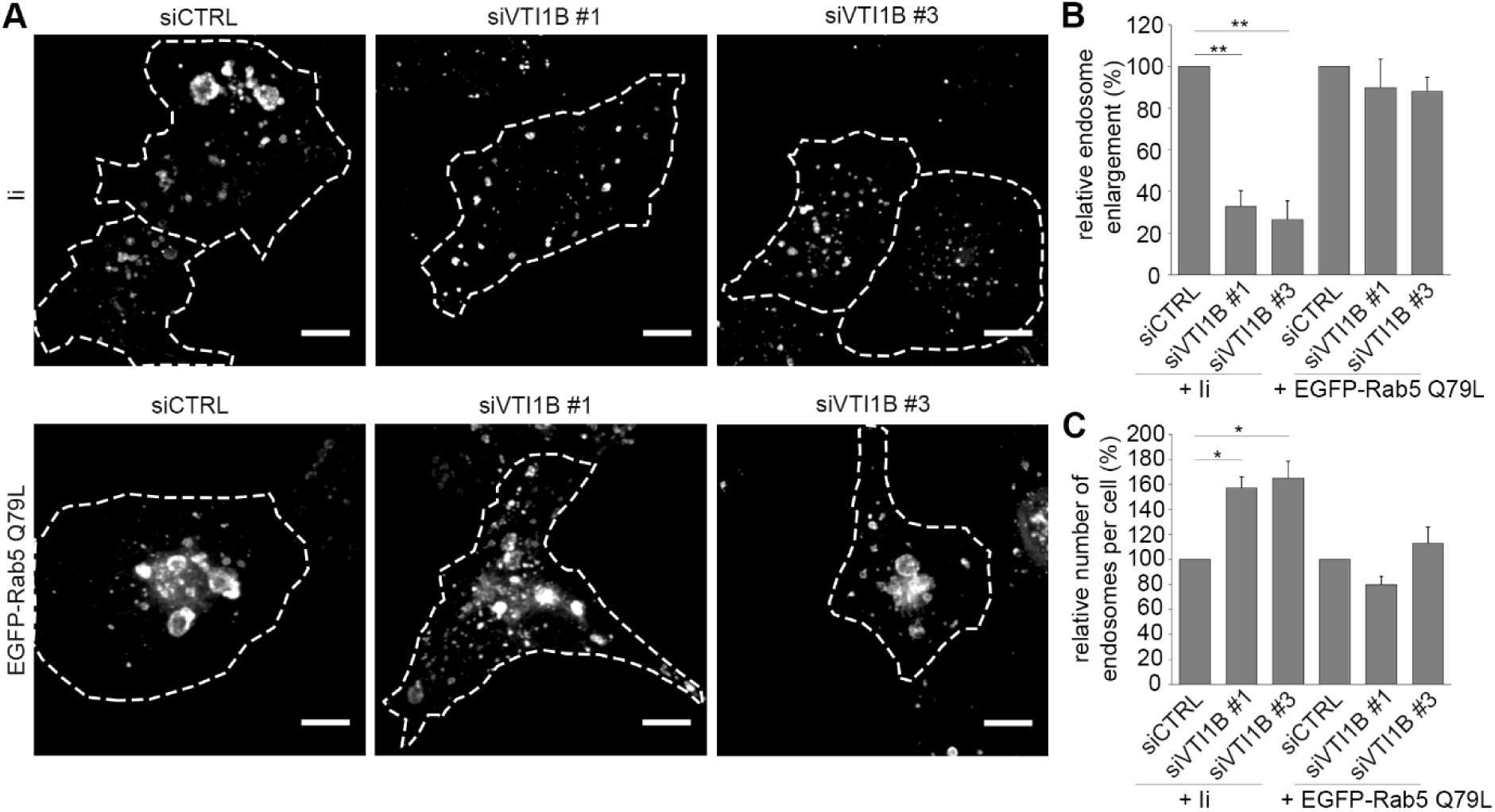
Vti1b specifically reduce Ii-mediated endosome enlargement. A) M1-pMep4-Ii cells were transfected with control siRNA (siCTRL) or siRNAs targeting VTI1B (siVTI1B #1 and siVTI1B #3). After 48h of transfection cells were either treated overnight with 7µM CdCl_2_ to induce Ii expression (top row) or transfected with EGFP-Rab5Q79L (bottom row) as indicated. Cells expressing Ii have been treated with anti-Ii antibody coupled with an Alexa Fluor 488 fluorescent dye for 30 minutes before fixation and imaging. Confocal images are shown. Dashed lines indicate the shape of the cells. Scale bars: 10 µm. B) Quantification of the percentage of endosomes (diameter ≥3 µm) in control and Vti1b silenced cells expressing either Ii or EGFP-Rab5Q79L. Data represent the mean ± s.e.m. of three different experiments (n=50). C) Quantification of the number of endosomes per cell in control and Vti1b silenced cells expressing either Ii or Rab5Q79L. Data represent the mean ± s.e.m. of three different experiments.

### Vti1b localized to contact sites between Ii-positive endosomes during their fusion

Given the well documented role of SNARE proteins in membrane fusion, we decided to evaluate the localization of Vti1b in M1 pMep4-Ii cells transfected with Ii and Vti1b-mCitrine. Interestingly, cells expressing Vti1b-mCitrine showed that this SNARE protein is localized at the contact sites between Ii-positive endosomes (Fig 4A, Movie EV1, see also 4B top row for fixed cells). In particular, Vti1b-mCitrine was detected directly at the opening fusion pore between two Ii-positive endosomes (Fig 4A, Movie EV1). Moreover, Vti1b-mCitrine was localized in intense microdomains on endosomes with these domains often found at the contact sites between two Ii-positive endosomes. As a control we used again mCherry-Rab5Q79L induced enlarged endosomes (Fig 4B bottom row). Quantification of the frequency with which Vti1b localized at the contact sites showed that Vti1b accumulated at about 80% of the contact sites between Ii-positive endosomes whereas Vti1b localized to this contact zone in 40% of the recorded interaction sites between Rab5Q79L-positive endosomes (Fig 4C). This shows that Vti1b is recruited more often to the contact site of fusing Ii-positive endosomes as compared to Rab5Q79L-positive endosomes (Fig 4C). Furthermore, we also evaluated the localization of endogenous Vti1b in M1 pMep4-Ii cells where Ii has been induced and endosomes were enlarged (Fig 4D). Similarly, to the transfected Vti1b-mCitrine fusion protein, immunofluorescence of endogenous Vti1b showed its localization at the contact sites of Ii-positive endosomes (Fig 4D). This was also studied in the Meljuso cells, and again Vti1b localized to contact sites between Ii-positive endosomes (Fig 4E), indicating that Vti1b associated with invariant chain at endosomal contact sites also in professional antigen presenting cells. In Meljuso cells the endosomes were not enlarged showing that Vti1b localization was not restricted to Ii induced enlarged endosomes.

**Figure 4.**
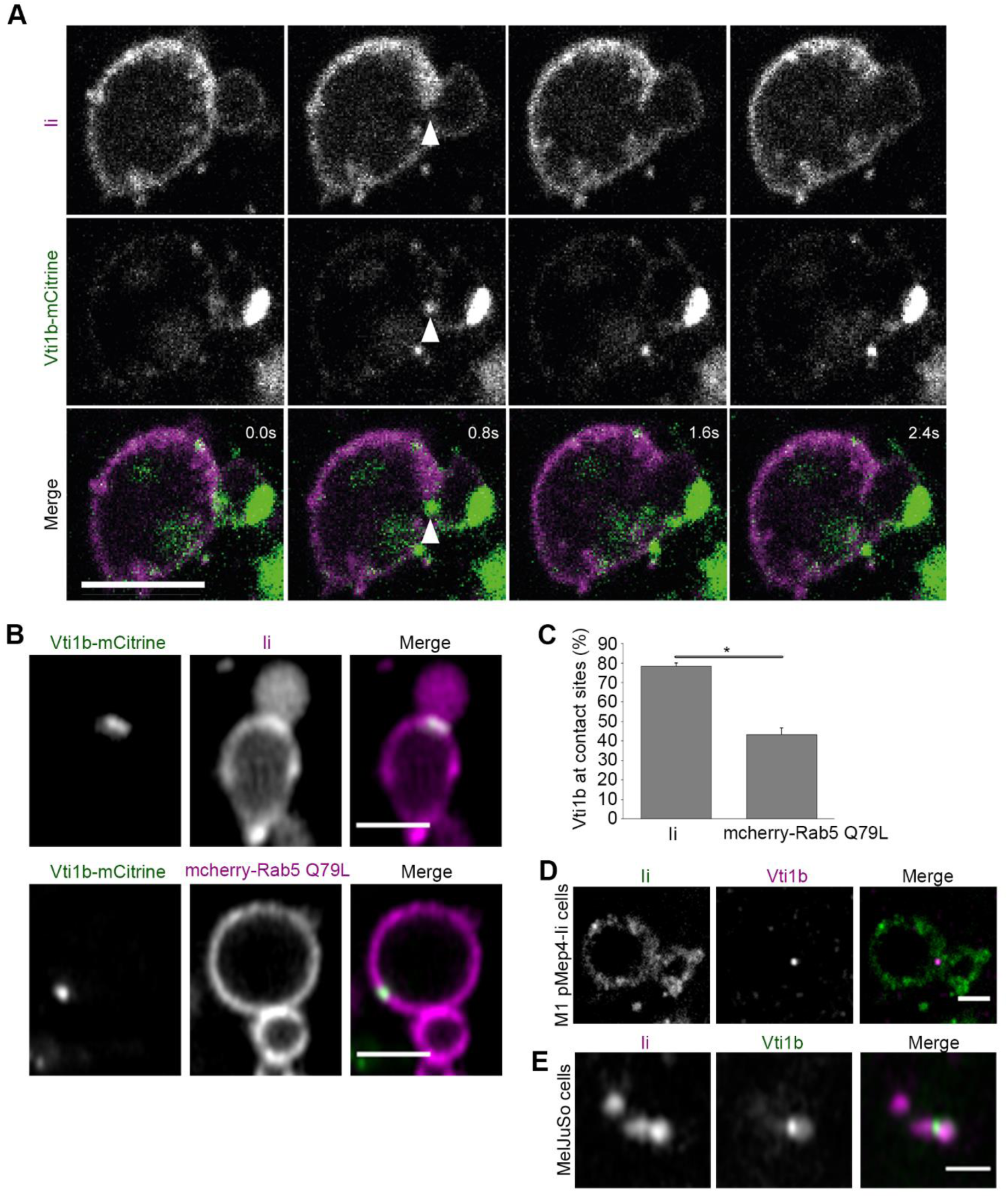
Vti1b localizes to contact sites at Ii-positive endosomes during fusion. A) M1 pMep4-Ii cells were transiently transfected with Vti1b-mCitrine o/n and Ii was expressed for 8 h (7µM CdCl_2_ added). Cells were treated with anti-Ii antibody coupled with Alexa Fluor 647 for 1 hour before live imaging. Time-lapse confocal images of Ii (magenta) and Vti1b-mCitrine (green) during endosome fusion are shown. Scale bar: 2 µm. B) M1 pMep4-Ii cells were either transiently transfected with Vti1b-mCitrine and treated with 7µM CdCl_2_ for 8h to induce Ii expression (top row) or co-transfected with Vti1b-mCitrine and mCherry-Rab5Q79L (bottom row). Cells expressing Ii were treated with anti-Ii antibody coupled with an Alexa Fluor 647 fluorescent dye antibody for 1 hour before imaged live by Fast AiryScan. Scale bar: 2 µm. C) Quantification of the percentage of Vti1b localized at the contact sites in cells expressing also Ii or Rab5Q79L as indicated is shown. Data represent the mean ± SEM of three independent experiments. D) M1 pMep4-Ii cells were treated with 7 µM CdCl_2_ for 8 h to induce Ii expression. Cells were treated with anti-Ii antibody coupled with an Alexa Fluor 647 fluorescent dye for 1 hour before fixation and immunostaining for Vti1b. Representative images of Ii (green) and endogenous Vti1b (magenta) positive endosomes are shown. Scale bars: 2 µm. E) Meljuso cells were treated with anti-Ii antibody coupled with an Alexa Fluor 647 fluorescent dye for 4 hours before fixation and immunostaining for Vti1b. Representative images of Ii (magenta) and endogenous Vti1b (green) positive endosomes are shown. Scale bar: 1 µm.

### Vti1b interacts with Ii

As Vti1b has a role in the regulation of the Ii-mediated endosomal enlargement and colocalize with Ii, we tested, whether this SNARE could interact with Ii. Firstly, we transfected either EGFP (as a negative control) or Vti1b-mCitrine into M1 pMep4-Ii cells. The expression of Ii was induced by CdCl_2_ treatment overnight and then we performed a co-IP experiment using anti-GFP nanobodies. We evaluated the presence of Ii in the eluates by Western blot (Fig 5A). We observed a band for Ii after immunoprecipitation of Vti1b-mCitrine with anti-GFP antibodies but not for the control EGFP (Fig 5A) showing that Vti1b interacts with Ii. In order to further elucidate the interaction between Vti1b and Ii, we transfected Vti1b-mCitrine and His-tagged Ii wt or Ii lacking the cytosolic tail (IiΔ27) in HeLa cells. As a negative control we used cells co-transfected with EGFP and Ii wt. Then we performed a co-immunoprecipitation assay with anti-GFP nanobodies (Fig 5B). Interestingly, Vti1b-mCitrine interacted with full-length Ii but also with IiΔ27 (Fig 5B), showing that the cytoplasmic tail of Ii is dispensable for binding to Vti1b.

**Figure 5.**
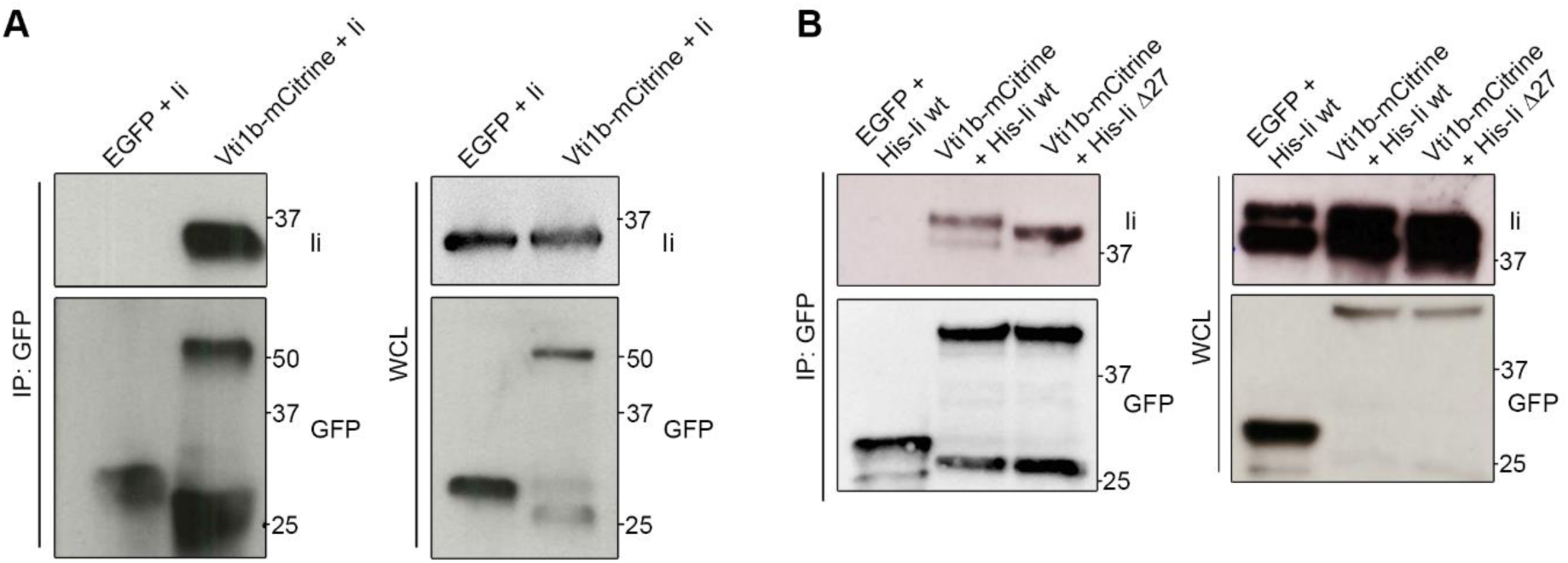
Vti1b interacts with Ii. A) M1-pMep4-Ii wt cells were transiently transfected with EGFP or Vti1b-mCitrine and Ii expression was induced by treatment with CdCl_2_ overnight. Cells were lysed and co-IP with GFP-Trap magnetic beads. Whole-cell lysates (WCL, on the right) and immunoprecipitated (IP, on the left) were subjected to Western blot analysis. B) HeLa cells were transiently co-transfected with either EGFP or Vti1b-mCitrine and Ii wt or mutated as indicated in the figure. Cells were lysed after 24 hours of transfection and thereafter co-IP with GFP-Trap magnetic beads. Whole-cell lysates (WCL, on the right) and immunoprecipitates (IP, on the left) were subjected to Western blot analysis. GFP and Ii was visualized using the corresponding anti-GFP or anti-Ii antibodies.

### Tailless Invariant chain relocates Vti1b to the plasma membrane

Vti1b contains a very short luminal domain (3 amino acids) and a long cytoplasmic tail (Brunger 2005) whereas Ii has a short (30 amino acids) cytoplasmic domain and a long luminal domain (Schroder 2016). Ii without a cytoplasmic tail is known to target invariant chain to the plasma membrane (PM) (Bakke and Dobberstein 1990).

To characterize the localization of these two associated proteins, we expressed the tailless Ii with a hexahistidine tag, (His-Ii Δ27) and Vti1b-mCitrine in HeLa cells and as control the same set up with full length Ii. As shown in Figure 6, Vti1b with full-length Ii was located in intracellular vesicles including endosomes positive for Ii. However, when Vti1b was expressed with Ii Δ27, a large fraction of Vti1b colocalized with Ii Δ27 at the PM. Since Vti1b was found to bind both Ii wild type and Ii Δ27, this suggest that these two molecules are associated in the biosynthetic pathway and that Vti1b is actively sorted by Ii to the endosomal pathway.

**Figure 6.**
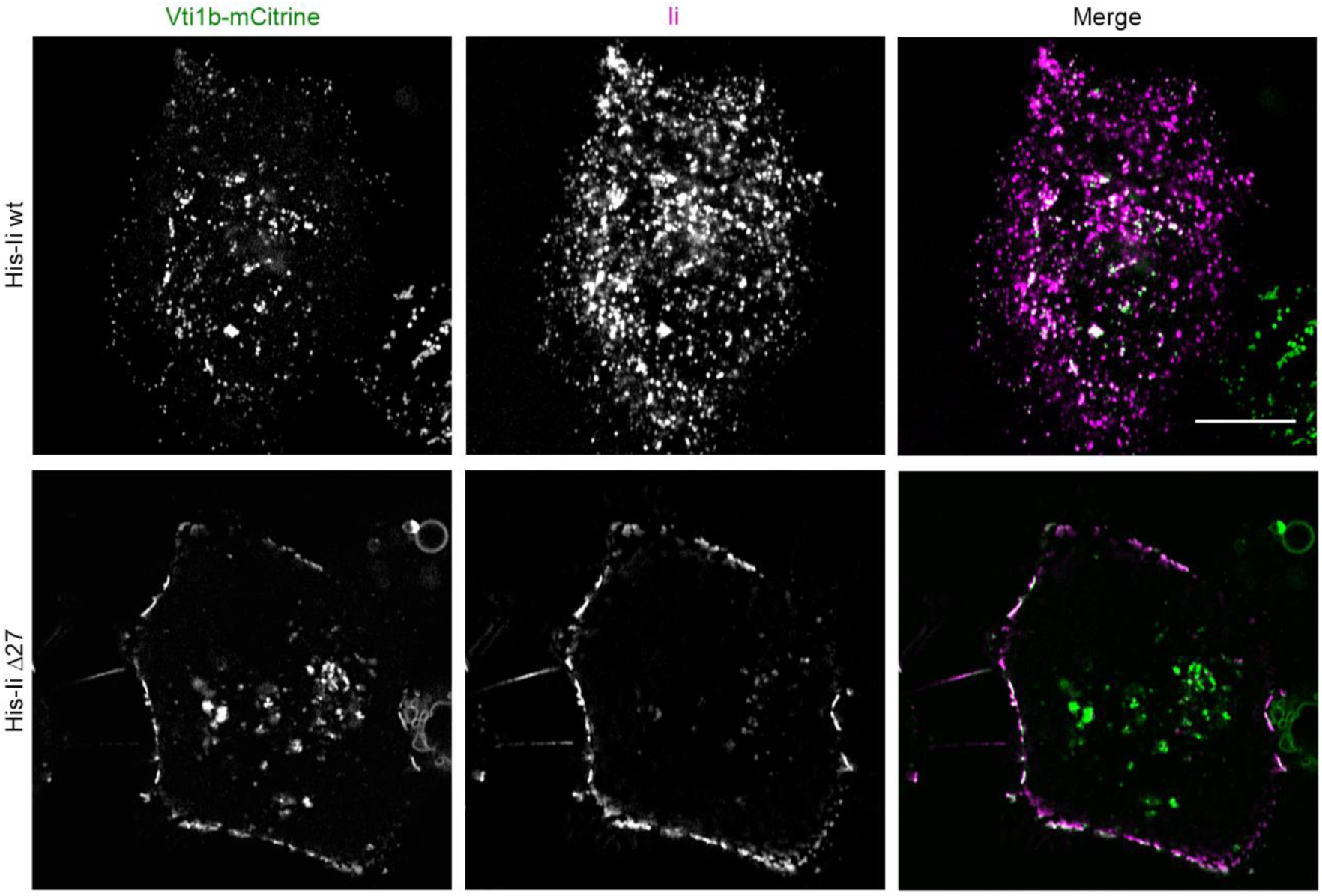
Vti1b and Ii Δ27 colocalize at the plasma membrane. HeLa cells were transfected with Vti1b-mCitrine and His-Ii (wt or Δ27) as indicated in the figure and then imaged. Confocal images are shown. Scale bar: 10 µm.

### The influence of Ii KO on Vti1b distribution in antigen presenting cells

In the intracellular trafficking experiments above we transfected Ii and mutants into cells lacking MHCII and associated genes (M1 and Hela cells). Meljuso cells, however, express endogenously the proteins involved in MHCII loading including MHCII, Ii and HLA-DM (Elliott and Neefjes 2006). MHCII may change the activities of Ii on endosomal maturation and expansion. To test this, we transfected Vti1b-mCitrine into Meljuso Ii KO and Meljuso wt cells followed by fixation and labelling with anti-CI-M6PR antibody that labels trans-Golgi network (TGN) and endosomes (Fig 7). Interestingly, Vti1b-mCitrine colocalized more with CI-M6PR in the Ii KO cells as compared with the wild type (Fig 7A, for quantitation, Figure 7B). We also evaluated the number of vesicles positive for Vti1b-mCitrine which was drastically reduced with about 40 vesicles/cell in the Ii KO cells as compared an average of 240 vesicles/cell for the wild type (Fig 7C). The volume of the CI-M6PR positive structures, in the Ii KO and wild type Meljuso cells were not significantly different (Fig 7D). As shown in Figure 7 we were able to rescue the phenotype by transfecting cDNA for Ii into the KO cells indicating that the altered intracellular distribution of Vti1b is indeed caused by the Ii KO and not by an off target effect.

**Figure 7.**
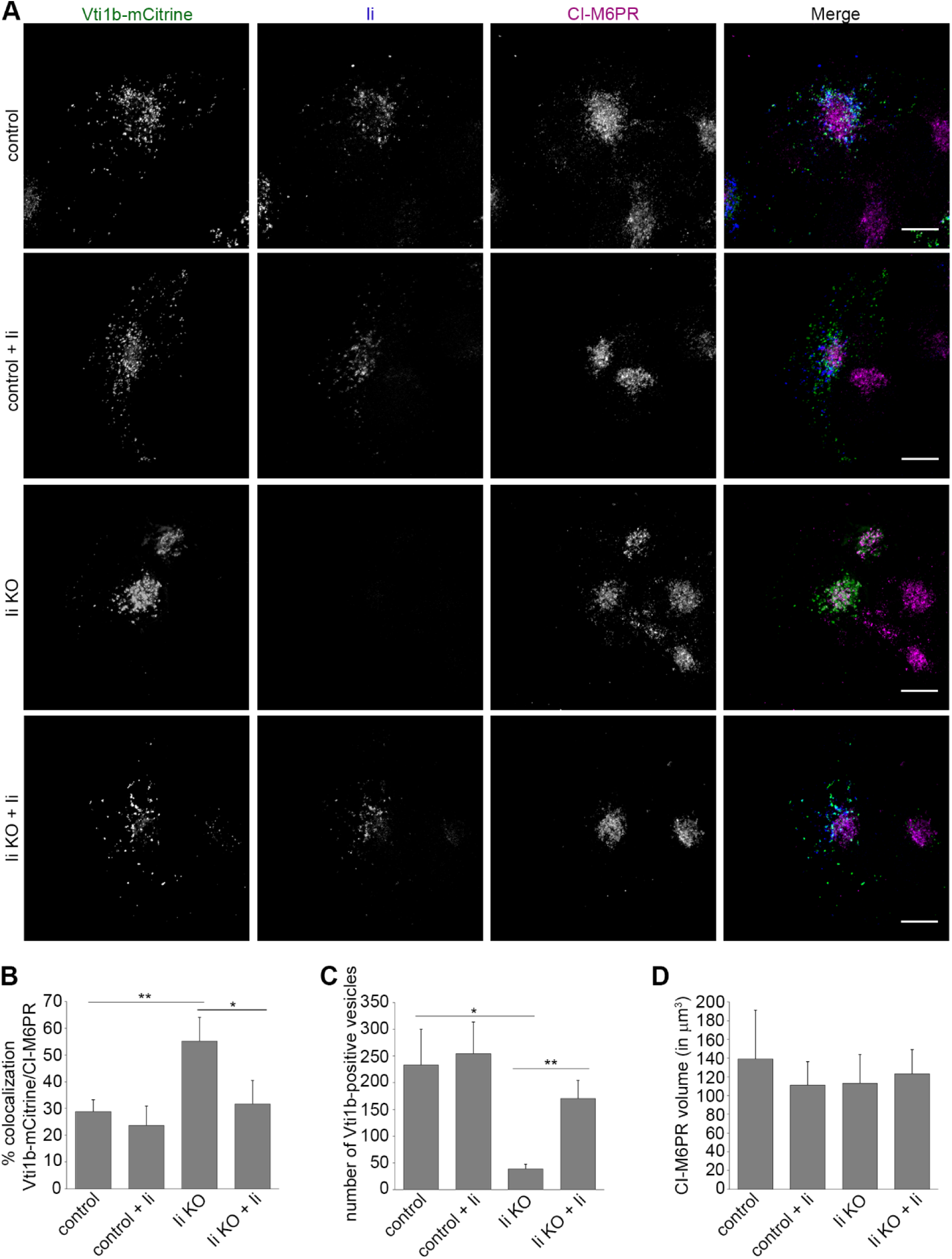
Vti1b localization is altered in Meljuso Ii KO cells. A) Meljuso control and Ii KO cells were transfected with either Vti1b-mCitrine alone or Vti1b-mCitrine and His-Ii as indicated in the figure and subsequently labelled live for Ii by treating the cells with an anti-Ii antibody coupled with an Alexa Fluor 647 for 4 h. Cells were stained after fixation with an anti-CI-M6PR antibody. Representative images (maximal projections) of Vti1b-mCitrine (green), Ii (blue), CI-M6PR (magenta) and merge are shown. Scale bars: 10 µm. B) Quantification of the percentage of colocalization between Vti1b-mCitrine and CI-M6PR in control, Ii-overexpressing, Ii KO or Ii-transfected Ii KO cells is shown. C) Quantification of the number of vesicles per cell positive for Vti1b-mCitrine in control, Ii-overexpressing, Ii KO or Ii-transfected Ii KO cells. D) Quantification of the volume of CI-M6PR positive structure in the indicated samples. Data represent the mean ± SEM of at least three independent experiments.

The CI-M6PR localizes in the trans-Golgi, plasma membrane and the late endosomal pathway (Hille-Rehfeld 1995). To firmly establish the localization of Vti1b in the Meljuso Ii KO, we also colocalized Vti1b with the Golgi resident protein giantin (Linstedt and Hauri 1993) and with TGN46 which mainly resides in the TGN (Prescott et al. 1997). As shown in Figure 8 more intense colocalization with both of these Golgi proteins was observed in the Ii KO cells.

**Figure 8.**
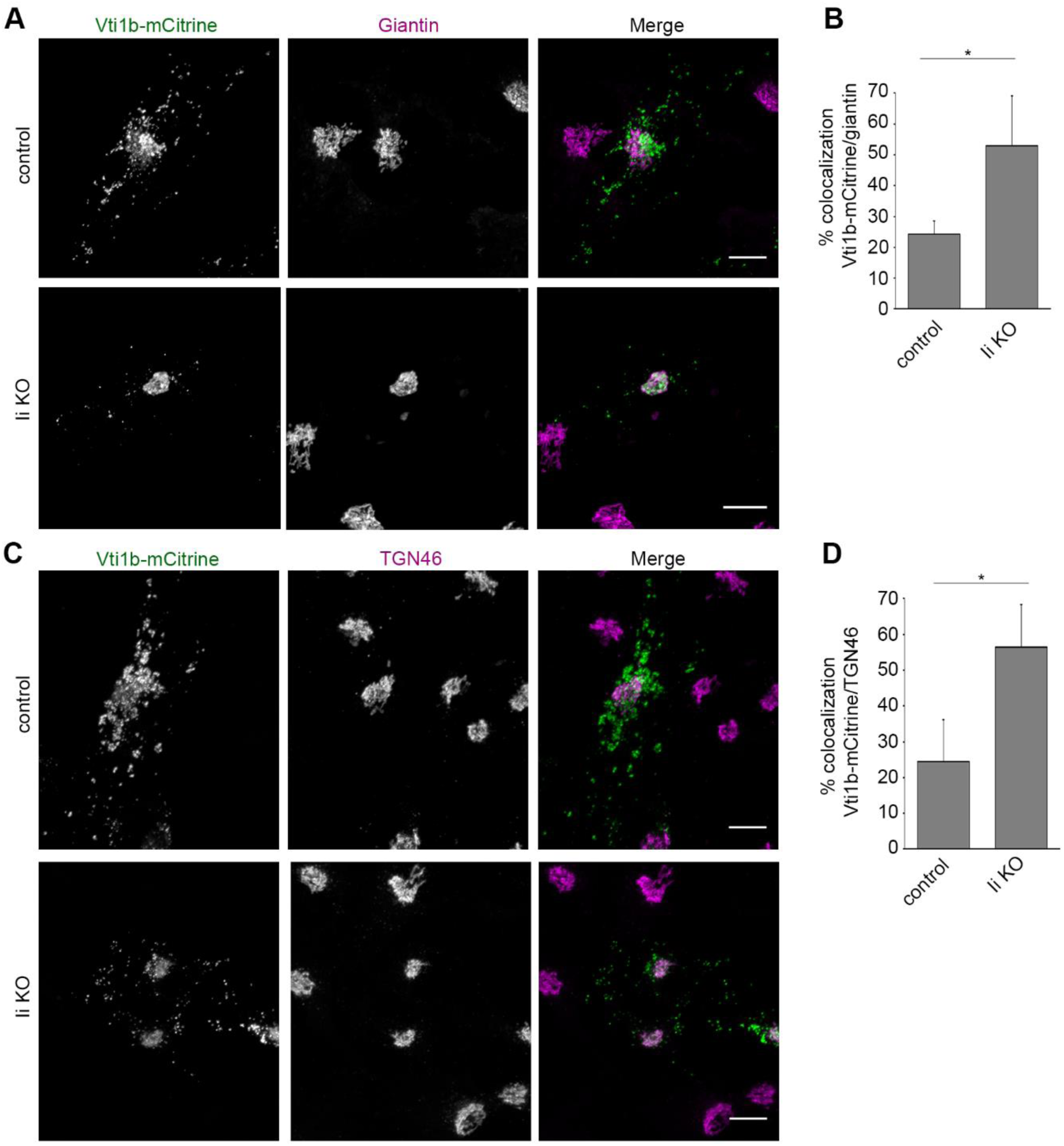
Vti1b is more localized on giantin and TGN46 positive structures in Meljuso Ii KO cells. A) Meljuso control and Ii KO cells were transfected with Vti1b-mCitrine and subsequently stained after fixation with an anti-giantin antibody. Representative images (maximal projections) of Vti1b-mCitrine (green), giantin (magenta) and merge are shown. Scale bars: 10 µm. B) Quantification of the percentage of colocalization between Vti1b-mCitrine and giantin in control and Ii KO cells is shown. C) Meljuso control and Ii KO cells were transfected with Vti1b-mCitrine and subsequently stained after fixation and permeabilization with an anti-TGN46 antibody. Representative images (maximal projections) of Vti1b-mCitrine (green), TGN46 (magenta) and merge are shown. Scale bars: 10 µm. B) Quantification of the percentage of colocalization between Vti1b-mCitrine and TGN46 in control and Ii KO cells is shown.

Vti1b has been reported to be in a complex with other endosomal SNARE proteins (Antonin et al. 2000; Wade et al. 2001; Pryor et al. 2004). However, the colocalization of CI-M6PR with fusion proteins of Vamp7, Stx8 and VAMP8 was not altered in the Ii KO cells (Fig EV1), indicating that the effect of Ii is not general for endosomal SNAREs and possibly specific for Vti1b.

### Vti1b is expressed in APCs and its protein levels are not affected by Ii expression

Given the important role of Ii in professional APCs and in antigen presentation (reviewed in (Schroder 2016)), we determined Vti1b protein level in APCs and other cell types. The expression levels of Vti1b in monocyte-derived dendritic cells, EBV immortalized B cells, the melanoma cell line MelJuSo, HeLa, Raji, M1 and Neuro2A cells was analysed by Western blot (Fig EV2A). Vti1b was widely expressed (Fig EV2A). To test whether Ii affects the expression levels of this SNARE protein, we analyzed Raji B cells (Fig EV2B,C) or the melanoma cell line MelJuSo (Fig EV2D,E) both wt and Ii KO. In both cell lines the expression of Vti1b was not affected by Ii expression. Similarly, high levels of Ii expression in M1 pMep4-Ii cells after induction with CdCl_2_ (Fig EV2F) did also not affect Vti1b expression (Fig EV2). Ii expression does not affect endogenous Vti1b expression.

### Knockdown of Vti1b does not affect lysosome dynamics and expression of endosomal marker proteins

Vti1b is reported to be a SNARE mainly involved in the late steps of endocytosis (Antonin et al. 2000; Pryor et al. 2004; Luzio et al. 2009; Kreykenbohm et al. 2002) and we wondered whether Vti1b knock down could affect the expression of other proteins relevant in the endosomal pathway (Rab7a, Rab7b, Rab9, LAMPI and EEA1, see Fig EV3). Western blot analysis revealed that the expression level of these endosomal proteins was unaltered after Vti1b depletion (Fig EV3B-F). In addition, we were also unable to detect significant alterations in the endosomal system in the Vti1b depleted M1 cells, even when Ii was expressed (Movie EV1).

### Silencing of Vti1b abrogates the endosomal maturation delay induced by Ii

Ii expression delays endosomal maturation from early Rab5 positive endosomes to late Rab7 positive endosomes (Gorvel et al. 1995; Landsverk et al. 2011) which correlates to Ii induced endosomal enlargement (Nordeng et al. 2002; Gregers et al. 2003). It would then be imperative that depletion of Vti1b would counteract the Ii induced endosomal maturation delay. The rate of endosomal maturation was determined by colocalization of the fluorescent fluid phase marker dextran-AlexaFluor 488 with Lysotracker Deep Red as a function of time in live cells. When we performed the assay in M1 pMep4-Ii cells without adding CdCl_2_ and thus not expressing Ii, 65% of dextran was localizing with Lysotracker Deep Red after 1 h. When expression of Ii was induced by CdCl_2_, entry of dextran was strongly delayed with only 35% overlap after one hour (Fig 9). This again involved Vti1b as silencing of Vti1b before induction of Ii expression restored the Ii delayed rate of endosomal transport and fusion to close to that of the control cells not expressing Ii. This suggests that Vti1b is essential for the Ii induced delay in endosomal maturation.

**Figure 9.**
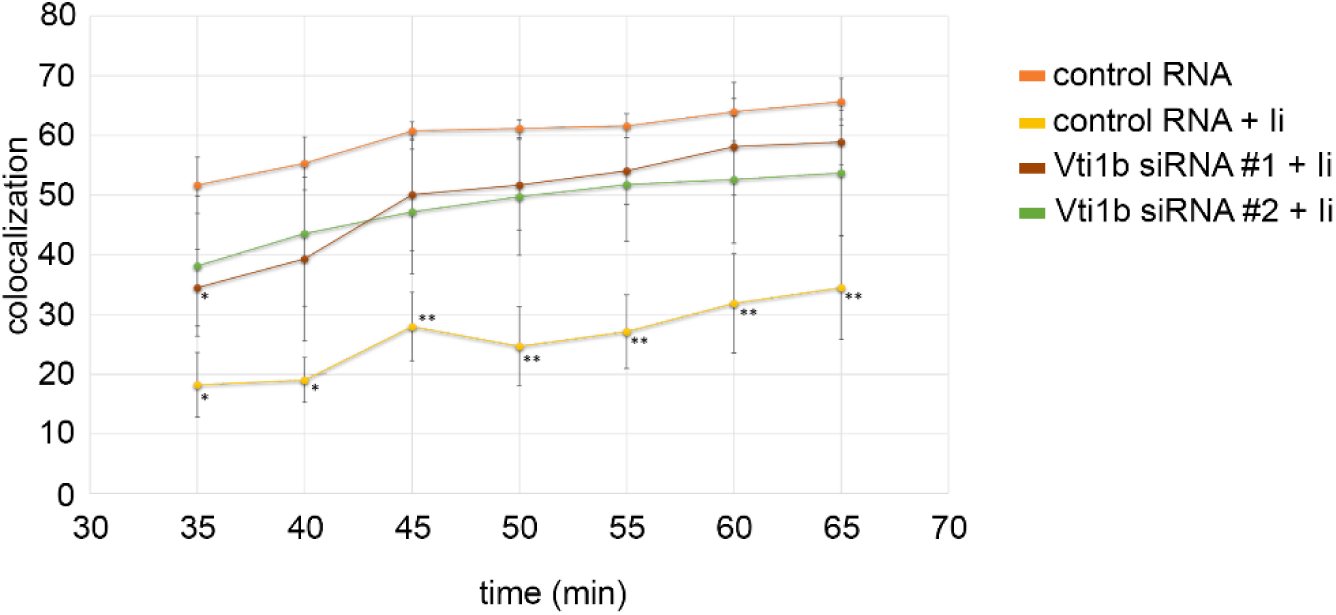
Ii causes a delay in trafficking dependent on Vti1b. Colocalization of dextran and Lysotracker Deep Red in M1-pMep4-Ii cells expressing or not Ii and treated with control siRNA or siRNAs against VTI1B as indicated.

## Discussion

Invariant chain associates with MHCII and the first function to be described was its ability to sort MHCII molecules to intracellular endosomal compartments. It was later shown that Ii actively sort other molecules to the endosomal pathway such as CD1 and MHCI, both molecules that participates in antigen presentation to specific T cells (for reviews see (Neefjes et al. 2011) and (Schroder 2016)). Other striking trafficking features of Ii is that, when expressed in non-antigen-presenting cells, it delays endosomal maturation (Gorvel et al. 1995; Landsverk et al. 2011). In addition, Ii has “fusogenic” properties as expression in cells lead to homotypic fusion of early endosomes (Nordeng et al. 2002; Romagnoli et al. 1993; Stang and Bakke 1997). This had led to the hypothesis that Ii is a key immunological molecule that is essential for creating “immunoendosomes” suitable for slowly processing endosomal contents for antigen loading (Neefjes et al. 2011). The MHCII-positive multivesicular late endosomal compartment in antigen-presenting cells (Peters et al. 1995) should then be a typical immunoendosome. The idea that Ii may delay endosomal maturation was until now only seen in fibroblast cells when transfected with Ii (Gorvel et al. 1995) which may fundamentally differ from professional antigen presenting cells. Here we show that KO of Ii in the antigen-presenting Raji B cell line and the MHCII positive Meljuso cells also affects endosomal maturation, in this case the Ii KO results in faster maturation. No direct correlation has been shown between the Ii induced formation of enlarged early endosomes and Ii induced maturation delay, however, mutations of the charged residues from negative to positive in the cytosolic tail of Ii has been found to abrogate both effects without altering the leucine sorting signals (Gregers et al. 2003; Nordeng et al. 2002) indicating that both effects are caused by similar mechanisms.

How would Ii manipulate the endosomal pathway? We have shown by NMR studies that the cytosolic tails of invariant chain could interact in trans and possibly interact in homotypic fusion (Motta et al. 1997). Ii has a short cytosolic tail of 30 aa, which would prevent it as acting as a genuine SNARE. It is thus more likely that other molecules are recruited to Ii to manipulate the endosomal pathway. We showed earlier that EEA1 or Rab5 were not essential for the Ii induced fusion and resulting enlargement of endosomes (Nordeng et al. 2002). However, the chemical compound NEM, that inhibits the SNARE function blocked fusion and formation of Ii induced enlarged endosomes (Nordeng et al. 2002), bringing our attention to SNAREs as essential for the process. We confirmed this data in the system that we used for the screen (Fig. 2B).

Based on these data we performed an RNAi screen to search for specific SNAREs that were essential for the Ii induced endosomal fusion. In our RNAi screen, we found that silencing of the Qb SNARE, Vti1b strongly decreased Ii-induced endosomal size (Fig 2C-E). Interestingly, although Vti1b has a major role in the fusion of late endosomes, it has not been implicated in the fusion of early endosomes (Luzio et al. 2009; Pryor et al. 2004; Kreykenbohm et al. 2002; Antonin et al. 2000). This shows that Ii employs a SNARE complex that is independent of the canonical early endosomal fusion machinery and can act in parallel.

In support of this, enlargement of endosomes induced by the expression of the constitutively active mutant of Rab5, Rab5Q79L, was not affected by Vti1b silencing (Fig 3). Interestingly, the reduction of Ii-positive endosomal size was accompanied by an increase in the number of Ii-positive endosomes, corroborating the hypothesis that Ii-induced enlargement of endosomes is due to an increase in endosomal fusion rather than reduced fission.

In support of this data, we demonstrated that both overexpressed and endogenous Vti1b was localized at the contact sites of Ii-positive endosomes in different cell lines and especially that Vti1b was directly at the opening fusion pore of fusing endosomes (Fig. 4; Movie EV1).

Strikingly, the effect was highly pronounced for the Ii-enlarged endosomes and Vti1b was detected at 80% of the endosome-endosome contact points. Using another method that enlarges endosomes, the constitutive active Rab5 (Rab5Q79L) (Wegner et al. 2010), Vti1b was only seen in 40% of the contact sites. This indicates that Vti1b is located at the site where fusion is happening and invariant chain expression then affects the location of Vti1b. Immunoprecipitation studies showed that Vti1b binds to Ii, directly or indirectly and this interaction may be the reason for the presence of Vti1b in the contact zone between Ii-positive endosomes. The fact that tailless Ii (Ii Δ27) also co-isolated Vti1b, suggests that the interaction is via their transmembrane domains or indirect.

The antigen presenting cell line Meljuso expresses Ii, and we find that the distribution of Vti1b was shifted perinuclear in the Ii KO cells showing an increased colocalization with CI-M6PR, Giantin and TGN46 (Fig. 7,8), which all are located mainly at the Golgi and correspondingly fewer Vti1b-positive vesicles were detected in Ii KO cells (Fig. 7). This suggest that Ii in wt Meljuso cells is involved in transport of Vti1b to the endosomal pathway.

Vti1b can move from TGN to late endosomes by binding to the AP1 interactor Clint1/EpsinR (Hirst et al. 2004). Ii, in addition to interacting to AP1, can also interact with AP2, responsible for internalization from the plasma membrane (Kongsvik et al. 2002; Hofmann et al. 1999). Ii is essential for efficient traffic of MHCII to the endosomal pathway and efficient antigen presentation (reviewed in (Neefjes et al. 2011)), but can also deliver other transmembrane proteins to endosomes, including CD70 (Zwart et al. 2010) and the FcR (Ye et al. 2008). We now suggest that Vti1b, due to its interaction with Ii, in antigen-presenting cells is sorted to the early endosomal pathway and thus participates in controlling endosomal maturation and fusion. As silencing of Vti1b leads to less endosomal fusion, we may also postulate that the Vti1b is instrumental for the Ii induced endosomal fusion (Nordeng et al. 2002; Romagnoli et al. 1993; Stang and Bakke 1997).

We here show that both the fusogenic properties of Ii and the maturation delay depend on the presence of Vti1b as silencing Vti1b corrected the Ii induced maturation delay in M1 cells. In this case, increased fusion mediated by Ii and the SNARE Vti1b and possibly other interactors on early endosomes would lead to a prolonged EE phase. Ii is also found to delay protein proteolysis (Gregers et al. 2003; Landsverk et al. 2011) and the part of the endosomal pathway that is trafficked by invariant chain would then create an ideal pathway for proteolytically processing instead of rapidly degrading antigenic proteins for MHC antigen loading of both MHCI, MHCII and CD1 that all associate with Ii.

Based on these data, we propose the following model (Fig 10). Ii binds directly or indirectly Vti1b for transport to endosomes, where can then engage in trans-SNARE complex formation. Higher degree of endosomal fusion dependent on a relocation of the “late” SNARE, Vti1b to early endosomes prolongs the EE phase and leads to a slower maturation creating a less proteolytic endosomal pathway. The model further postulate that as Ii positive endosomal vesicles engage in homotypic fusion, then not only Ii but also the associated molecules like MHCII will be concentrated in some late endosomal/lysosomal vesicles as in the MIIC compartment (Peters et al. 1991).

**Figure 10.**
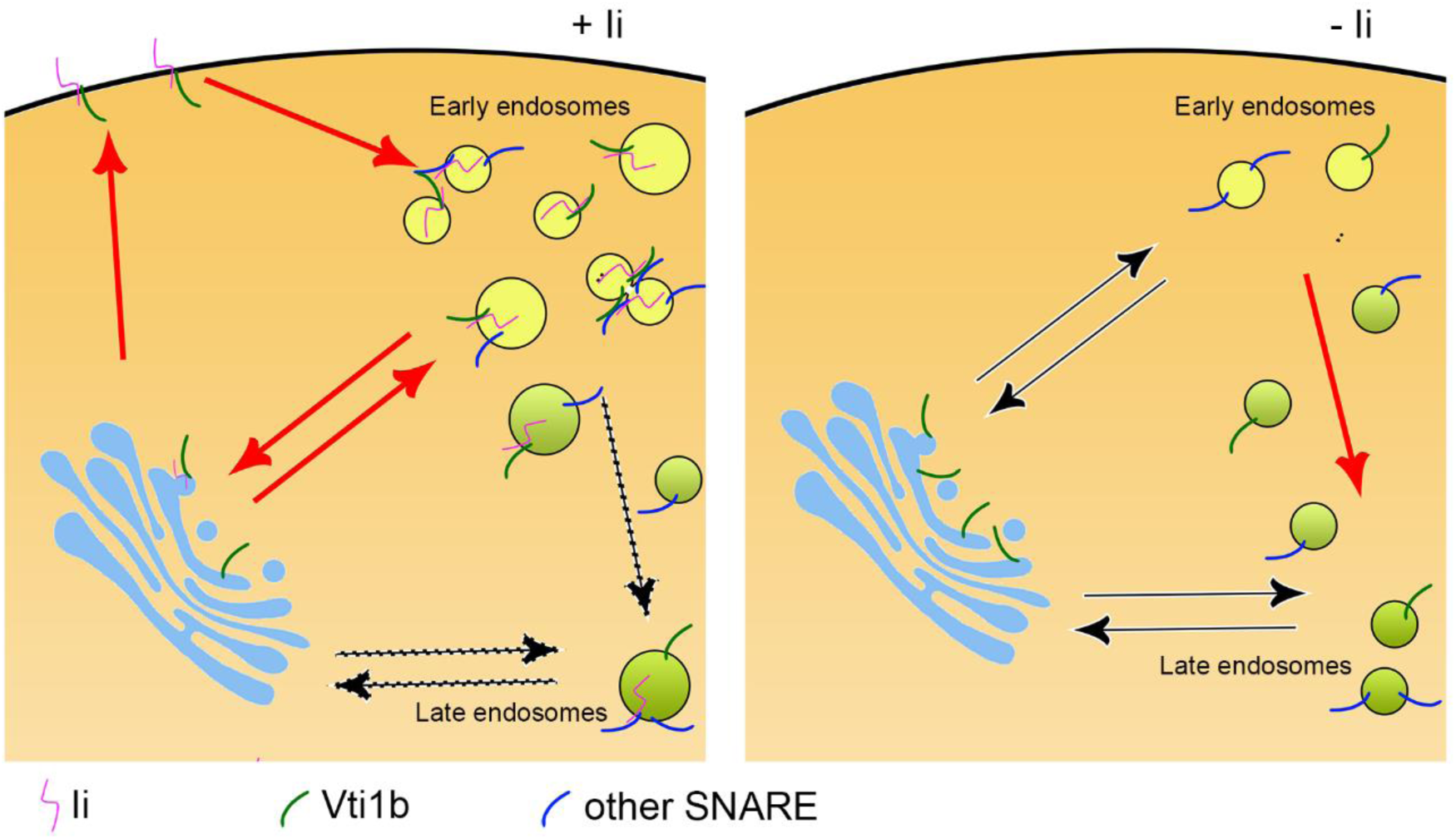
Schematic model depicting the main findings of this study. Ii binds to Vti1b and facilitates its trafficking via the Golgi apparatus to the plasma membrane and from there to early endosomes. There, Vti1b forms a SNARE complex leading to membrane fusion. Increased endosomal fusion dependent on Vti1b delays endosomal maturation (left). Lack of Ii induces a reduction of the endosomal localization of Vti1b, less endosomal fusion and faster endosomal maturation (right).

It is very interesting that the expression of a single protein, in this case Ii, a protein specific to antigen-presenting cells has the ability to reshape the endosomal pathway in a specific way. Our results support the notion that Ii not only increases the fusion of early endosomes by employing the late endosomal Qb-SNARE Vti1b but also affects endosomal maturation. Therefore, Ii is not only chaperoning MHCII to the loading compartment (MIIC), but is also involved in shaping this specialized organelle of professional APCs. The MIIC has a unique proteolytic environment, less aggressive than that of classical lysosomes (Neefjes 1999) which is needed for efficient but not full processing of antigens. We suggest that this property is intricately connected to Ii’s ability of delaying endosomal maturation. In addition, the antigen loading compartments integrate antigen input from several other sources than endocytosis, such as phagocytosis, macropinocytosis and autophagy (Blum, Wearsch, and Cresswell 2013), but whether the transfer of these to the immunoendosome is by a general or a more specific mechanism is not known.

In conclusion, we show here that the Qb SNARE Vti1b is involved in Ii-mediated endosomal fusion. Ii redistributes Vti1b from the Golgi to early endosomes, which leads to increased fusion and also delayed maturation of endosomes. This mechanism of regulating endosomal fusion could be critical for efficient MHCI and II antigen presentation and thus normal immune responses.

## Materials and Methods

### Cell lines

The human fibroblast-like M1 cells (Royer-Pokora, Peterson, and Haseltine 1984) stably transfected with pMep4-Ii (Skjeldal et al. 2012), the human melanoma cell line Meljuso (Johnson et al. 1981), the human lymphoblast cell line Raji (Pulvertaft 1964), the human adenocarcinoma HeLa cells (ATCC CCL-2) and the rat neuronal Neuro2A cell line (Cogli et al. 2013) were described before. Healthy donor derived Epstein-Barr virus growth-transformed lymphoblastoid cell lines (EBV-LCL) were provided by Sebastien Wälchli. Mononuclear cells were isolated from buffy-coats from healthy donors through density-gradient centrifugation by using Lymphoprep® (Axis Shield). Buffy-coats from anonymous donors were obtained from the local blood bank (Section for Immunology and Blood Transfusion, Ullevål University Hospital, Oslo, Norway) according to the guidelines of the local blood bank approved by the Norwegian Regional Committee for Medical Research Ethics. Monocyte-derived dendritic cells (MDDCs) were generated from plastic adherent or directly isolated monocytes (Monocyte Isolation Kit II, Miltenyi Biotec) through culture for 6 days in Roswell Park Memorial Institute medium (RPMI-1640; (Lonza) containing 100 ng/ml granulocyte-macrophage colony stimulating factor (GM-CSF; Immunotools) and 20 ng/ml IL-4 (Invitrogen, Life Technologies) supplemented with 10% fetal calf serum (FCS), 2 mM L-glutamine, 100 U/ml penicillin and 100 µg/ml streptomycin (Sigma).

M1 and HeLa cell lines were grown in Dulbecco’s Modified Eagle’s Medium (DMEM; Lonza) supplemented with 10% FCS, 2 mM L-glutamine, 100 U/ml penicillin and 100 µg/ml streptomycin (all Sigma). Additionally, 200 µg/ml hygromycin B (Sigma) was added to stably transfected M1 cells. Meljuso, Raji, EBV and MDDCs were grown in RPMI-1640 supplemented with 10% FCS, 2 mM L-glutamine, 100 U/ml penicillin and 100 µg/ml streptomycin.

For live-cell microscopy, cells were grown in 35mm glass bottom dishes (MatTek), µ-Slide 8 Well glass bottom (ibid), or 4-chamber 35mm glass bottom dish (CellVis) for 24 h in complete medium.

### Establishment of CRISPR/Cas9 Ii KO cell lines

Raji Cas9 and Ii knockout cell lines to be described elsewhere. Meljuso cells were transfected with px330 Sp-Cas9 and sgRNA vectors (12456 and 12458 targeting Ii exon 1) together with a vector coding for blasticidin resistance using Lipofectamine® 2000. Successfully transfected cells were selected by treatment with blasticidin for 1 week. Colonies grown from resistant cells were picked, expanded and screened for Ii knockout by Western blot.

### Constructs

mEGFP-C1 was a gift from Michael Davidson (Addgene plasmid # 54759). His6-Ii p33 wt and His6-Ii Δ27 in pCGFP-EU were purchased from GenScript. pCGFP-EU empty vector has been described before (Kawate and Gouaux 2006). mcherry-Rab5 Q79L has been described before (Bergeland et al. 2008). The Vti1b-mCitrine plasmid was constructed as follows: the coding sequence of human VTI1B was amplified by PCR using 5’ and 3’ primers containing an EcoRI and a BamHI restriction site, respectively. Forward primer was: 5’-AATCAGAATTCATGGCCTCCTCCG-3’. Reverse primer was 5’- CATCGGATCCCAATGGCTGCGAAAGAATTTG-3’. The fragment was then subcloned into pECFP-N1 plasmid cut with EcoRI and BamHI. The identity of all plasmids was confirmed by Sanger sequencing (GATC biotech).

### Antibodies and reagents

Primary antibodies used were: mouse anti-Ii (clone M-B741, targeting the extracellular/luminal domain, 555538, 1:500) and mouse anti-EEA1 (610456, 1:200 for immunofluorescence; 1:1000 for western blot) from BD Biosciences; mouse anti-tubulin (13-8000, 1:12000) from Life Technologies; rabbit anti-Rab7a (2094S, 1:500) from Cell Signaling; rabbit anti-GFP (ab6556, 1:2000), rabbit anti-Vti1b (ab184170, 1:200 for Western blot, 1:100 for immunofluorescence), rabbit anti-CI-M6PR (ab32815, 1:200) and mouse anti-Rab9 (ab2810, 1:300) from Abcam; mouse anti-Rab7b (H00338382-M01, 1:300) from Abnova, and mouse anti-LAMPI (H4A3, 1:500 for immunofluorescence; 1:4000 for western blot) from Developmental Hybridoma Bank. AlexaFluor™ secondary antibodies (Invitrogen) were used for immunofluorescence analysis at 1:200 dilution. Secondary antibodies conjugated with horseradish peroxidase (GE Healthcare) were used at 1:5000 for immunoblotting.

Anti-Ii for live immunostaining was labelled using Monoclonal antibody labeling kit (Molecular Probes) according to manufacturer’s protocol with AlexaFluor™-488, −555, or - 647, respectively.

Hoechst dye (H3569, Life Technologies) was used at 0.2 µg/ml. LysoTracker® Red DND-99 (LTR), used at a concentration of 100 nM, LysoTracker® DeepRed, used at a concentration of 50 nM, 8-Hydroxypyrene-1,3,6-trisulfonic acid trisodium salt (HPTS), used at 1 mM concentration, and dextran AlexaFluor 488 10,000 MW, used at a concentration of 0,5 mg/ml, were purchased from Thermo Fisher.

Poly-L-lysine (PLL, 0.01% solution), Wortmannin and N-ethylmaleimide (NEM) were all purchased from Sigma.

### siRNA transfection

Individual siRNAs were transfected by forward transfection using Lipofectamine® RNAiMax reagent (Thermo Fisher) according to manufacturer’s protocol. In brief, 250.000 cells/well were seeded in 6-well plates (VWR) in 1 ml antibiotic-free medium 24h prior to transfection. Transfection mixes were prepared by diluting 2.5 µl siRNA stock (80 µM) in 250 µl Opti-MEM and 6 µl RNAiMax in 250 µl Opti-MEM followed by immediate mixing of both. The transfection mixture was then incubated at room temperature for 20 min before addition to the cells. 4h after transfection 1 ml of complete medium was added. Medium on the cells was replaced the next day. 48h after transfections, cells were washed twice with pre-warmed PBS, trypsinized for 1min, resuspended in 2 ml complete medium and split between 35 mm imaging dishes (MatTek) or 8-well slides (ibidi) for imaging and 6-well plates for lysis and subsequent biochemical assays.

### Plasmid transfection

For plasmid transfection, cells were transfected using Lipofectamine® 2000 (Thermo Fisher) according to manufacturer’s protocol. In brief, cells were seeded the day before transfection. Before transfection, the medium was replaced by Opti-MEM® (Thermo Fisher). Transfection mixes were prepared with 1.5 µl Lipofectamine® 2000 (Thermo Fisher) per 1 µg of plasmid DNA. The transfection mix was then added drop-wise to the cells.

### Immunofluorescence

For immunofluorescence, cells were grown on 10×10mm #1.5 high precision coverslips (Marienfeld). Cells were washed twice with ice-cold PBS, fixed at room temperature for 15 min with 3% PFA, quenched for 2 min with 50 mM NH4Cl, washed again twice with PBS, permeabilized for 15 min with 0.25% saponin (Sigma) in PBS, incubated with primary antibody for 20 min, washed 3 times for 5 min each with 0.1% saponin in PBS, incubated with appropriate secondary antibody for 20 min in the dark, washed three times for 5 min each with 0.1% saponin in PBS, washed twice with PBS, and mounted with mowiol on glass slides (Thermo Scientific).

### Endocytic trafficking assay

Cells were grown for 24h on 35mm glass bottom dishes (MatTek) or 35 mm 4-chamber glass bottom dishes (CellVis) in complete medium without phenol red. LTR was added and cells incubated for another 15 min at 37°C before addition of HPTS, 10 min before start of time-lapse imaging. For dextran internalization assay, cells have been incubated with 0,5 mg/ml dextran and Lysotracker Deep Red for five minutes. Then, cells have been washed three times and imaged every five minutes.

### Confocal Imaging

For the siRNA screen, imaging was performed on an Olympus FV1000 XV81 confocal laser scanning microscope (inverted) equipped with a UPLSAPO 60×1.35NA oil immersion objective. Images of cells transfected with individual siRNAs were acquired on a Leica SP8 DMi8-CS confocal microscope equipped with a HC PL Apo 40x 1.3NA oil immersion objective.

Live imaging of transfected cells and imaging of immunostained cells was performed on a Zeiss LSM880 microscope equipped with a Plan Apo 63x 1.4NA oil immersion objective. Live cells were imaged using Fast AiryScan mode with optimal sampling for super-resolution.

Time-lapse imaging to study the trafficking of HPTS to acidic compartments was performed on a Zeiss LSM880 microscope, equipped with a Plan Apo 40x 1.3NA oil immersion objective. Z stacks of 5 slices each were acquired at intervals of 1 µm every 10 min for a total of 200 min at multiple positions. Hardware-based autofocusing (DefiniteFocus.2) was performed at every position and every time-point. Fluorescence emission of HPTS and LTR was separated by linear unmixing based on reference spectra obtained from single stained samples.

### Image processing and analysis

Endosome size was quantified by using ImageJ software. In brief, confocal 3D stacks were maximum projected, in case of EGFP-Rab5Q79L images, the cytosolic background was subtracted by rolling ball background subtraction. The resulting images where then binarized by using a manually adjusted threshold for each image. After that, the images were inverted, morphologically opened, objects split by watershed processing and manual splitting where needed. Finally, the analyse particles function was used to measure all objects. The Feret’s diameter was used as the measurement of object size and objects larger than 3 µm were considered enlarged endosomes.

LTR colocalization was quantified using the colocalization function in ZEN blue software (Zeiss). In brief, scatter plots were used to define thresholds on the centre z slice for each position. The ratio of overlapped pixels to all LTR-positive pixels was then calculated for all slices of a z stack, summed for each position and compared between conditions using excel software (Microsoft).

### Immunoblotting

Cell lysates were subjected to SDS-PAGE followed by blotting onto polyvinylidene fluoride (PVDF) membranes (Millipore), and probed with each primary antibody diluted in 2% blotting-grade non-fat dry milk (BioRad). Next, membranes were incubated with secondary antibodies conjugated to HRP and subsequently with SuperSignal West Femto Maximum Sensitivity Substrate (Thermo Fisher) before digital imaging (Kodak image station 4000R) or Amersham™ ECL Prime Western Blotting Detection Reagent before exposure on Amersham™ Hyperfilm ECL (both GE Healthcare).

### Statistical analysis

If not indicated otherwise, columns and error bars represent the mean ± standard error of the mean (s.e.m.) of at least three independent experiments. Treatments were compared to control using two-tailed Student’s t-test, either paired or unpaired as appropriate (excel). Differences were considered statistically significant at p<0.05 (indicated by *), p<0.01 (**) and p<0.001 (***), respectively.

### Co-Immunoprecipitation

Co-Immunoprecipitation experiments using GFP-trap® (ChromoTek) were performed according to manufacturer’s protocol. In brief, 1-2×10^6^ cells were seeded on 10 cm cell culture dishes and incubated for 24 h at 37°C, 5% CO_2_. The cells were then transfected using Lipofectamine 2000 as described earlier. M1 pMep4-Ii cells were treated with 7 µM CdCl_2_ overnight to induce expression of Ii. Cells were washed twice with ice-cold PBS and lysed in 200 µl lysis buffer (100 mM NaCl, 1 mM EDTA, Complete® protease inhibitors (Sigma), 1 mM PMSF (Phenylmethylsulfonylfluorid, Sigma), 0.5% NP-40) for 30 min on ice. Lysates were cleared by centrifugation at 13,000 x g for 10 min at 4°C, diluted with 300 µl buffer and loaded on magnetic agarose anti-GFP beads (GFP-trap®_MA, ChromoTek) for 1 h at 4°C. Beads were then washed three times. Proteins were eluted using 2x Laemmli sample buffer with freshly added D-thiotreitol (200 mM, Sigma) for 10 min at 95°C.

## Acknowledgements

We thank Sebastien Wälchli (University of Oslo) for providing the EBV cells and Cecilia Bucci (University of Salento) for providing the Neuro2A cells. We thank Michael Davidson (Florida State University) for providing the mEGFP-C1plasmid through Addgene and Jens P. Morth (University of Oslo) for the pCGFP-EU empty vector. Finally, we thank the NorMIC Oslo imaging platform, Department of Biosciences, University of Oslo.

We gratefully acknowledge the support from the Research Council of Norway (Grants 214183, and 230770 to OB) and the Norwegian Cancer Society (grant 113472 to OB).

## Author contributions

DF and AM designed, performed, analyzed the majority of the experiments, MD contributed to the IP experiments and WB experiment in different cell types. LJ and JN generated the Meljuso Ii knock-out cell line, MM and JT generated the Raji Ii knock-out cell line. OB coordinated the project and DF, AM and OB wrote the manuscript. All co-authors approved the final version of the manuscript.

