## Supplemental data for "Invariant chain regulates endosomal fusion and maturation through the SNARE Vti1b"

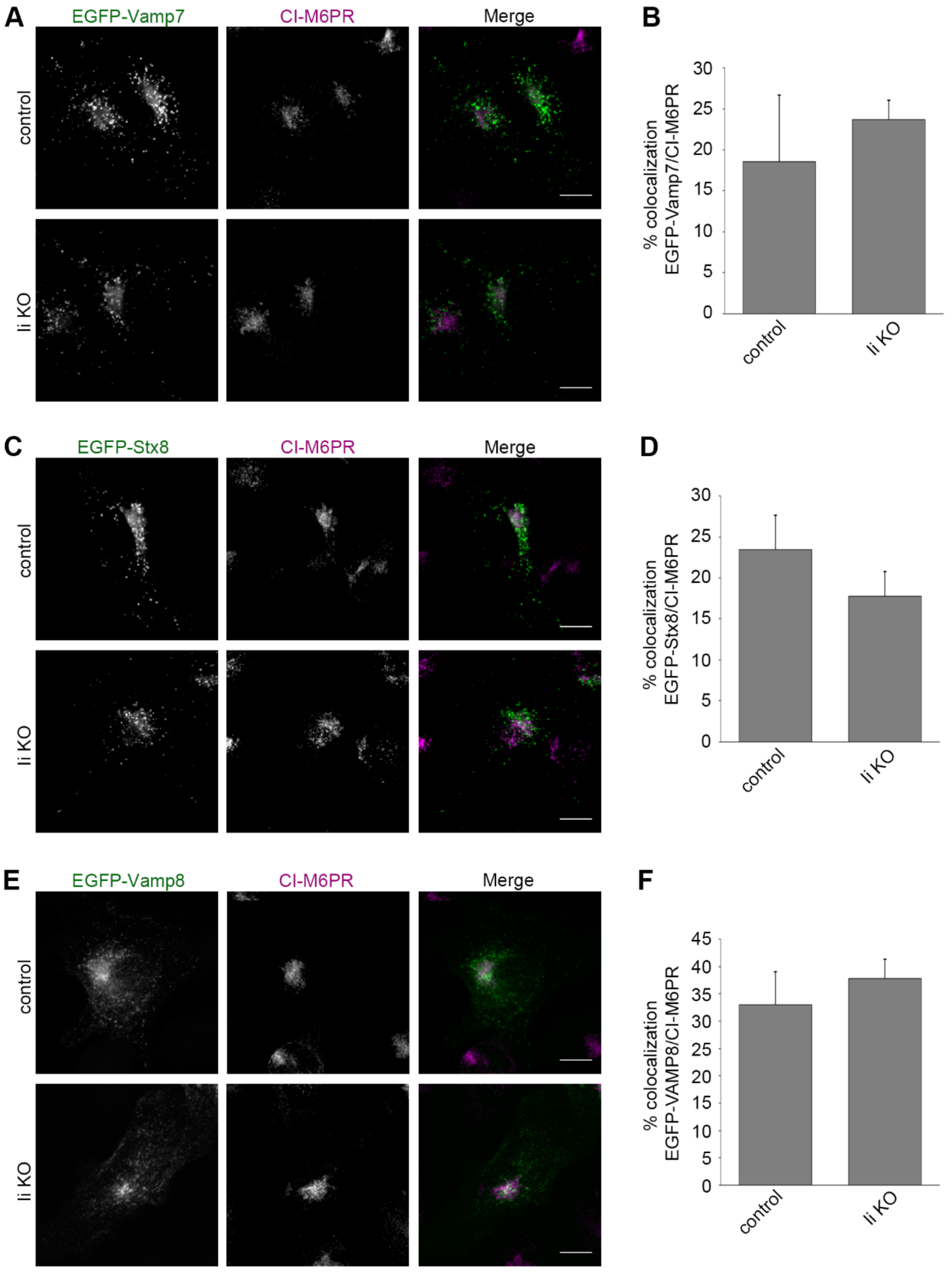


Figure EV1

**Figure EV1. Localization of Vamp7, Stx8 and Vamp8 is not altered in Meljuso Ii KO cells.** A) Meljuso control and Ii KO cells have been transfected with EGFP-Vamp7 and subsequently stained after fixation with an anti-CI-M6PR antibody. Representative images (maximal projections) of EGFP-Vamp7 (green) and CI-M6PR (magenta) and merge are shown. Scale bars: 10 µm. B) Quantification of the percentage of colocalization between EGFP-Vamp7 and CI-M6PR in control and Ii KO cells is shown. C) Meljuso control and Ii KO cells have been transfected with EGFP-Stx8 and subsequently stained after fixation with an anti-CI-M6PR antibody. Representative images (maximal projections) of EGFP-Stx8 (green) and CI-M6PR (magenta) and merge are shown. Scale bars: 10 µm. D) Quantification of the percentage of colocalization between EGFP-Stx8 and CI-M6PR in control and Ii KO cells is shown. E) MeljusO control and Ii KO cells have been transfected with EGFP-Vamp8 and subsequently stained after fixation with an anti-CI-M6PR antibody. Representative images (maximal projections) of EGFP-Vamp8 (green) and CI-M6PR (magenta) and merge are shown. Scale bars: 10 µm. D) Quantification of the percentage of colocalization between EGFP-Vamp8 and CI-M6PR in control and Ii KO cells is shown.

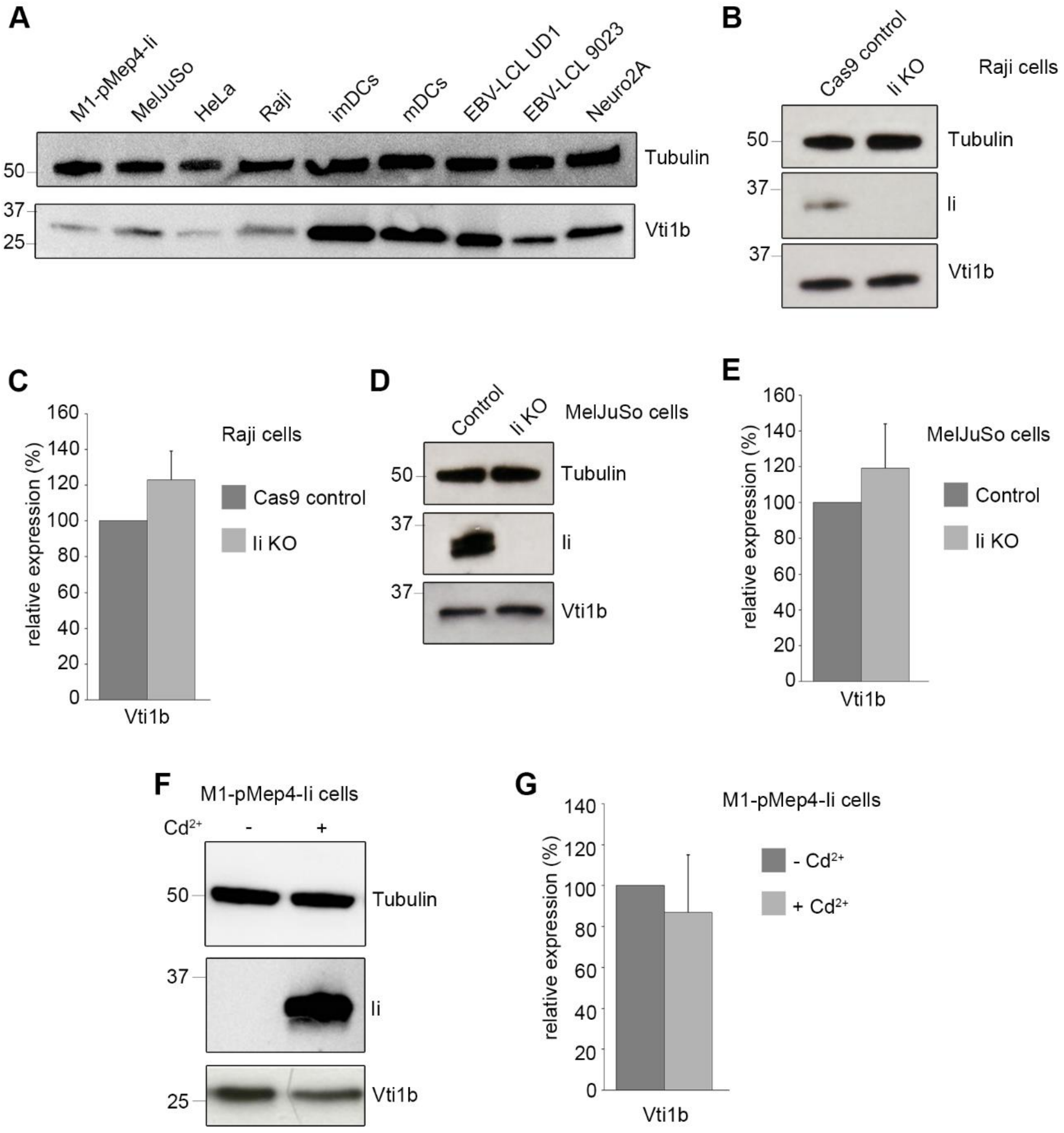


Figure EV2

**Figure EV2. Expression levels of candidate proteins in different cell types.** A) Lysates of several cell types as indicated in the figure have been subjected to western blot analysis using anti-Vti1b and anti-tubulin antibodies. B) Raji Cas9 control and Ii KO cells have been lysed and relative samples have been subjected to western blot analysis. Antibodies against Vti1b, Ii and tubulin have been used to detect their respective abundance. Tubulin has been used as loading control. C) Quantification of Vti1b abundance in Raji Cas9 control and Ii KO cells is shown. Data represent the mean ± s.e.m. of three independent experiments. D) Meljuso control and Ii KO cells have been lysed and relative samples have been subjected to western blot analysis. Antibodies against Vti1b, Ii and tubulin have been used. E) Quantification of Vti1b abundance in Meljuso control and Ii KO cells. Data represent the mean ± s.e.m. of three independent experiments. F) Control M1-pMep4-Ii wt cells not expressing Ii (-$Cd^{2+}$) or after treatment with 7µM $CdCl_2$ overnight to induce the expression of Ii (+$Cd^{2+}$) have been lysed. Lysates were subjected to western blot analysis using anti-Vti1b, anti-Ii and anti-tubulin antibodies. E) Quantification of Vti1b abundance in M1-pMep4-Ii wt cells expressing (+$Cd^{2+}$) or not (-$Cd^{2+}$) Ii is shown. Data represent the mean ± s.e.m. of three independent experiments.

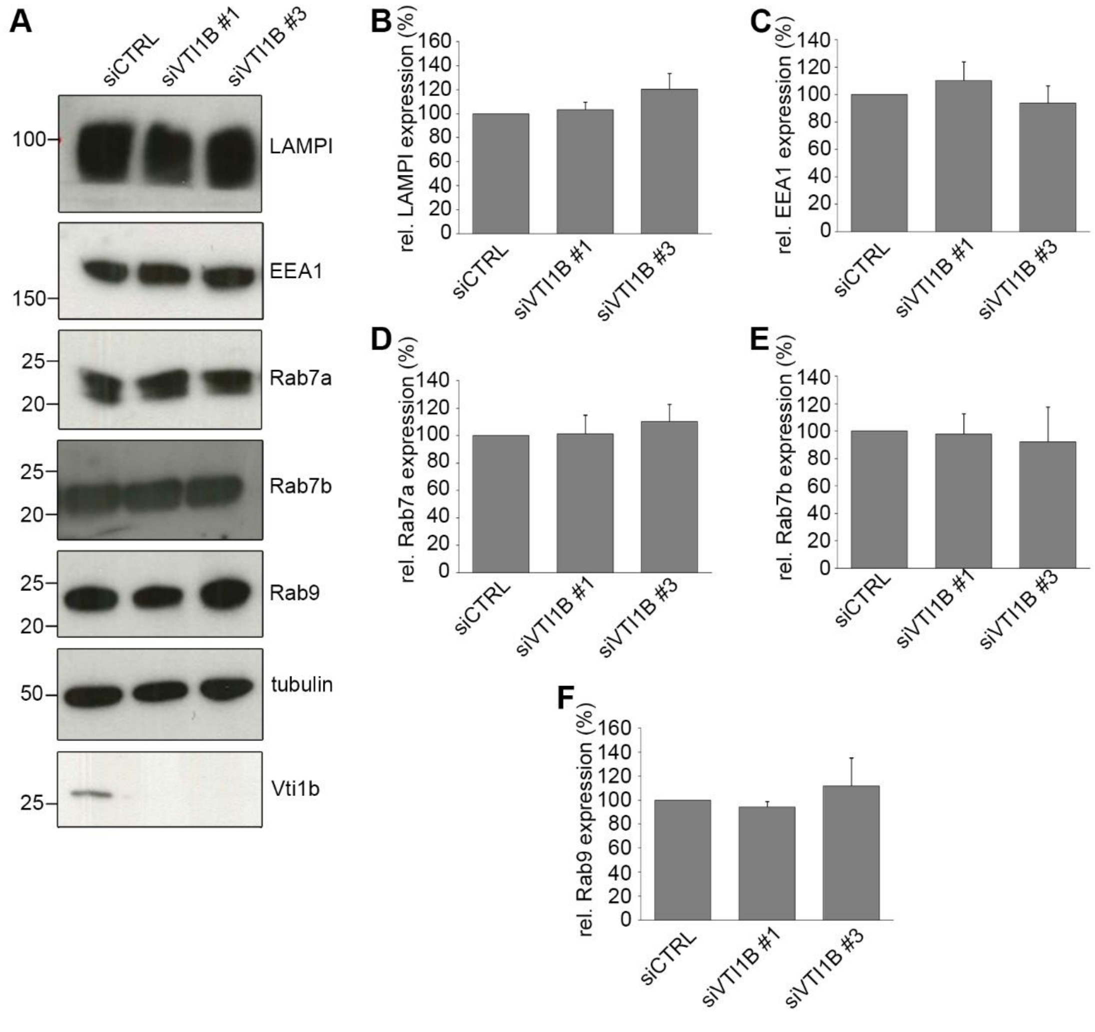


Figure EV3

**Figure EV3. Depletion of Vti1b does not alter expression of endosomal trafficking related proteins**. A) M1-pMep4-Ii cells have been transfected with either control or VTI1B siRNAs and subsequently lysed. Samples have been subjected to western blot analysis by using antibodies against LAMP1, EEA1, Rab7a, Rab7b, Rab9, Vti1b and tubulin. B) Quantification of LAMP1 abundance in control or Vti1b depleted cells. C) Quantification of EEA1 abundance in control or Vti1b depleted cells. D) Quantification of Rab7a abundance in control or Vti1b depleted cells. E) Quantification of Rab7b abundance in control or Vti1b depleted cells. F) Quantification of Rab9 abundance in control or Vti1b depleted cells. Data represent the mean ± s.e.m. of three independent experiments.

### Movie legends

**Movie EV1. Vti1b is localized to contact sites during fusion of Ii-positive endosomes.** M1 pMep4-Ii cells were transiently transfected with Vti1b-mCitrine and Ii expression was induced for 8 h with 7µM $CdCl_2$ treatment. Ii and Vti1b-mCitrine are indicated in magenta and in green, respectively. Scale bar: 2 µm.

**Movie EV2. Ii expression does not induce any effect dependent on Vti1b on dynamics of late compartments.** M1-pMep4-Ii cells were transfected with either control RNA or VTI1B siRNA and Ii expression was induced with 4h treatment with $CdCl_2$.. Internalization of anti-Ii antibody (tagged with fluorescent tag, green) and lysotracker DeepRed (red) has been conducted for 30 minutes. Cells were imaged with an Olympus Fluoview 1000 IX81 inverted confocal laser scanning microscope. Scale bars: 10 µm.
